# Isolation Strategy Shapes the Matrisome Landscape of Cancer-Associated Fibroblast Extracellular Vesicles

**DOI:** 10.64898/2026.05.22.727064

**Authors:** Mahmoud K. Eldahshoury, Emily Moss, Jacob Gillett-Woodley, Matthew S Hindle, Martha Ilett, Mark O. Collins, James R. Boyne

## Abstract

Cancer-associated fibroblasts (CAFs) secrete small extracellular vesicles (sEVs) that mediate stromal remodelling, tumour progression, and pre-metastatic niche formation. A foundational assumption in EV research is that MISEV-compliant preparations from the same conditioned medium are biologically equivalent. Here, we directly challenged this assumption through a side-by-side comparison of ultracentrifugation (UC), size exclusion chromatography (SEC), and the EXODUS nanofiltration platform using breast CAF-conditioned media, characterised in accordance with MISEV2023 guidelines using nanoparticle tracking analysis, cryogenic transmission electron microscopy (Cryo-TEM), and quantitative proteomics. EXODUS and SEC recovered approximately 7-fold more particles per mL than UC. While Cryo-TEM confirmed intact vesicle morphology across all methods, UC preparations exhibited substantial non-vesicular background, with gene ontology analysis revealing significant enrichment of ribosomal, mitochondrial, and ER-derived proteins absent from EXODUS and SEC. Matrisome profiling further uncovered method-dependent divergence in the composition of core versus matrisome-associated proteins, highlighting differences extending beyond standard purity metrics. These findings demonstrate that MISEV2023 compliance is necessary but insufficient for methodological equivalence. The isolation method should be treated as a biological variable and selected according to the EV subpopulation or cargo class under investigation.

## Introduction

The extracellular vesicle (EV) field faces a systemic challenge regarding reproducibility, particularly concerning the impact of isolation methodology on downstream biological interpretation (Rai *et al*., 2025; Morales-Sanfrutos *et al*., 2026). It is now widely recognised that isolating small extracellular vesicles (sEVs) from similar conditioned media using two different methods can recover preparations with fundamentally different protein compositions, particle concentrations, and functional activity profiles despite satisfying current minimal reporting standards (Lundy *et al*., 2026; Preußer *et al*., 2026). This discrepancy represents a significant analytical confound rather than a minor technical variation. When the isolation procedure determines the resulting proteome, methodological choice effectively masquerades as biological reality. Consequently, findings attributed to specific disease states, experimental conditions, or cell types may instead reflect the sedimentation kinetics or filtration properties inherent to the isolation workflow (Van Deun *et al*., 2014; Kowal *et al*., 2016; Jeppesen *et al*., 2019). This issue is most consequential when the source secretome is compositionally complex, such as in the tumour microenvironment (TME), where cancer-associated fibroblasts (CAFs) present precisely this scenario (Liu *et al*., 2026)

CAFs are found in solid tumours and are the principal architects of the tumour extracellular matrix (ECM) (Lu *et al*., 2025; Lupsa *et al*., 2026). Unlike the secretomes of standard immortalised cell lines, the CAF secretome is structurally dominated by large, insoluble, surface-associated ECM macromolecules, including collagens, proteoglycans, laminin heterotrimers, fibronectin, and a suite of matrix-remodelling enzymes that form a dense and physicochemically complex environment, making EV isolation from the CAF secretome particularly challenging (Belhabib *et al*., 2021; Yuan *et al*., 2023; Xie *et al*., 2025). Specifically, CAF-derived sEVs are increasingly recognised as direct vectors of this stromal programme, transferring ECM components, matrix metalloproteinases, and pro-fibrotic signals that promote tumour progression, therapy resistance, and pre-metastatic niche formation (Clément *et al*., 2022; Liu *et al*., 2023; Santi *et al*., 2024). The matrisome cargo of these vesicles is not incidental. It represents the definitive molecular signature of the CAF phenotype (Hynes and Naba, 2012; Sarkar *et al*., 2023). However, matrisome proteins are structurally complex, spanning a wide range of molecular weights, glycosylation states, and supramolecular assembly forms, properties that render them uniquely susceptible to differential recovery or loss depending on the isolation strategy employed (Mouw, Ou and Weaver, 2014). Therefore, the CAF secretome serves as an ideal and exceptionally demanding model to test whether the selection of an isolation strategy constitutes a fundamental biological variable.

Critically, compliance with the Minimal Information for Studies of Extracellular Vesicles (MISEV) framework does not resolve this methodological divergence. MISEV2023 compliance establishes minimum identity criteria to define EVs, such as the presence of transmembrane tetraspanins and cytosolic proteins, but it does not guarantee molecular equivalence between preparations from the same biological source (Théry *et al*., 2018; Welsh *et al*., 2024). A preparation that preserves the EV corona despite meeting minimal reporting standards yields a fundamentally different functional profile than one in which these critical surface interactions have been disrupted (Preußer *et al*., 2026).

This challenge is further exacerbated by the inherent heterogeneity of the CAF compartment. CAFs encompass distinct subpopulations, including myofibroblastic, inflammatory, and ECM-CAF, each with divergent secretory programmes and matrisome outputs (Fan *et al*., 2025; Wu *et al*., 2026). In this context, method-dependent differences in matrisome cargo recovery are not merely technical artefacts; they risk misrepresenting which CAF subpopulation’s biological programme is captured, with direct consequences for cross-laboratory comparability and biological interpretation. Despite the significance of this variable, no systematic comparison has evaluated isolation method performance specifically in the context of the CAF secretome.

Here, we report a controlled, side-by-side comparison of different isolation methods, including ultracentrifugation (UC), size-exclusion chromatography (SEC), and EXODUS for the isolation of EVs from primary human breast cancer-associated fibroblast (BCAF)-conditioned media. Using dual-platform nanoparticle tracking analysis (NTA), Cryo-TEM, MISEV2023-compliant characterisation, quantitative label-free proteomics, gene ontology enrichment analysis, and full matrisome annotation against the Naba classification framework. We demonstrate that these three methods recover a common EV pool; however, they enrich for biologically distinct populations of the CAF secretome from the same conditioned media. The differences are most pronounced and consequential at the level of ECM cargo. Our findings establish isolation method as a biological variable in CAF-EV research and argue that the choice of isolation strategy for ECM-rich stromal secretomes determines not only preparation quality but also the feasibility of the research questions being asked.

## Methods

### Collection of breast CAF-conditioned medium

Primary human breast CAF were obtained from the Breast Cancer Now tissue bank. CAF cells were grown and maintained with DMEM/F12 + 10% foetal bovine serum (FBS) (Gibco). The medium was replaced with a fresh, pre-warmed medium every 3 days. To collect CAF-conditioned media (CM) for EV isolation, cells were grown to 80% confluency and then washed with phosphate-buffered saline (PBS). CAF-CM was collected after 48 hrs of serum starvation in DMEM (Gibco, ThermoFisher) and subsequently centrifuged at 300 × g for 5 min to remove dead cells, then stored at −80°C for future use.

### Small EVs isolation

Conditioned medium was collected from cultured cells and centrifuged at 2,000 × *g* for 30 min using an Allegra X-30 centrifuge to remove cells and debris. The supernatant was passed through a 0.22 µm aPES membrane filter (Nalgene) and subsequently divided into three equal aliquots for isolation by UC, SEC, or the EXODUS H600 system. For UC-based isolation, samples were centrifuged at 100,000 × *g* for 90 min at 4°C using an Avanti JXN-26 ultracentrifuge with a JA-20 rotor. The resulting pellets were resuspended in 0.1 µm-filtered PBS and subjected to a second ultracentrifugation step at 100,000 × *g* for 60 min at 4°C using a Beckman Coulter Optima MAX-XP with a TLA-110 rotor. Final pellets were resuspended in 0.1 µm-filtered PBS. For SEC isolation, filtered conditioned medium was concentrated to 0.5 mL using a 10 kDa molecular weight cut-off protein concentrator (Vivaspin, Cytiva). The concentrate was applied to a qEVoriginal/35 nm Gen 2 column according to the manufacturer’s instructions. Fractions 7–11, previously identified as enriched in EVs with minimal protein background, were pooled and reconcentrated using 10 kDa MWCO spin concentrators (Shelton *et al*., 2025). For isolation using the EXODUS H600, samples were processed with the exosome isolation device (EID) unit size medium following the manufacturer’s protocol. After isolation, the central collection chamber was retrieved. Experimental details of the isolation procedure were submitted to EV-TRACK under the ID EV260037 (EV-TRACK Consortium *et al*., 2017)

### Nanoparticle tracking analysis

NTA was performed using a NanoSight NS300 (Malvern Panalytical) equipped with a 488 nm laser and a scientific CMOS camera. Isolated sEV were diluted in 0.1 µm filtered PBS to achieve a working concentration within the recommended detection range (1 × 10^6^ to 1 × 10C particles/mL). For each sample, three 90-second videos were recorded under constant camera settings. The flow mode syringe pump was set to level 50 to maintain a steady particle movement throughout acquisition. All measurements were performed at room temperature with the following settings: camera level 14 with a detection threshold of 5. Particle size distribution and concentration were determined based on analysis of particle Brownian motion using Malvern NS3000 software.

### ZetaView nanoparticle tracking

EV samples were diluted in 0.1 μm filtered ultrapure water to a final volume of 1 mL before analysis. Each measurement consisted of three cycles, with the instrument scanning 11 cell positions per cycle and capturing 60 frames per position. All measurements were conducted at a controlled temperature of 25°C. Data were analysed using ZetaView Software version 8.06.01 SP1.

### Western blot

Samples were lysed in radioimmunoprecipitation assay (RIPA) buffer supplemented with 1× protease and phosphatase inhibitor cocktail (Sigma, 20-188) and incubated on ice for 30 min. EV samples were spiked with recombinant enhanced green fluorescent protein (eGFP; Proteintech, EGFP) for normalisation. Total protein concentration was determined using a micro bicinchoninic acid (microBCA) assay (ThermoFisher, 23235) according to the manufacturer’s protocol, and all samples were normalised accordingly. Equal amounts of protein were prepared in reduced Laemmli sample buffer (6x), heated at 95°C for 5 min, and resolved via sodium dodecyl sulfate-polyacrylamide gel electrophoresis (SDS-PAGE). Proteins were transferred onto polyvinylidene difluoride (PVDF) membranes, which were subsequently blocked in 5% non-fat dry milk diluted in Tris-buffered saline with 0.1% (v/v) Tween-20 (TBST) overnight at 4°C. Membranes were incubated with primary antibodies against CD63 (Abcam, ab217345; 1:1000), CD81 (Proteintech, 66866-1; 1:1000), ALIX (Proteintech, 12422-1; 1:1000), and Calnexin (Proteintech, 10427-2; 1:2000) diluted in 3% non-fat dry milk in TBST and incubated overnight. After three washes in TBST, membranes were probed with the corresponding HRP-conjugated secondary antibodies: rabbit (phytoAB, PHY6000; 1:10000) or mouse (Sigma-Aldrich,61-6520; 1:5000) for 1 hour at room temperature. Protein bands were visualised using chemiluminescent substrate (Bio-Rad, 1705061) and visualised using a ChemiDoc Imaging System (Bio-Rad).

### Mass Spectrometry-based Proteomics

12.5 µl of each EV sample was mixed with an equal volume of 2X S-trap lysis buffer to a final concentration of 5% SDS/50 mM Tris, pH 7.4. Samples were reduced with TCEP to a final concentration of 5 mM and heated at 70 °C for 15 minutes. Proteins were alkylated by adding Iodoacetamide (IAA) to a final concentration of 10 mM, and the samples were incubated at 37 °C in the dark for 30 minutes. 2.5 µl of 12 % phosphoric acid and 330 µl of S-Trap binding buffer (90% methanol, 100 mM TEAB, pH 7.1) were then added to each sample. Samples were then loaded into S-Trap columns (ProtiFi) by centrifugation at 10,000 x g for 60 seconds. Samples were washed four times with 225 µl of S-Trap binding buffer through centrifugation at 10,000 x g for 60 seconds. 1 µg of Trypsin (Pierce, sequencing grade) was added to each sample in a total digestion volume of 20 µl of 50 mM TEAB, and digestion was performed at 47 °C for 1 hour, followed by 37 °C for 1 hour. Peptides were eluted with the addition of 40 µl of 50 mM TEAB, 40 µl of 0.2% aqueous formic acid, and then 40 µl of 50% acetonitrile with 0.2% formic acid, followed by centrifugation at 10,000 x g for 60 seconds for each elution. Pooled eluted peptides were dried in a vacuum concentrator and resuspended in 0.5% formic acid for LC-MS/MS analysis.

Each sample was analysed using nanoflow LC-MS/MS using a timsTOF FLEX MALDI-2 mass spectrometer equipped with a Captive-Spray II source coupled to a nanoELUTE LC System (Bruker). Peptides were desalted online using a Pepmap C18 nano trap column (300 μm I.D. x 5 mm, Thermo Fisher), then separated using a Pepsep 25cm C18 column (75 μm ID, 1.9 μm particles, Bruker) over a 30-minute gradient. The timsTOF FLEX MALDI-2 was operated in positive mode using a dia-PASEF method. The source parameters were as follows: capillary voltage, 1,600 V; dry gas, 3.0 L/min; and dry temperature, 180 °C. The MS1 and MS2 spectra were collected in the m/z range of 100–1,700. The collision energy was set by linear interpolation between 59 eV at an inverse reduced mobility (1/K0) of 1.60 versus/cm2 and 20 eV at 0.6 versus/cm2. DIA-PASEF acquisition was performed using a 1/K0 range of 0.6–1.6 and an m/z range of 400–1,200.

Raw mass spectrometry data files were processed with DIA-NN version 2.3.0. Data were analysed in library-free mode using a peptide library predicted from a Human proteome Fasta file (downloaded Aug 2025) containing 20475 proteins. The following settings were used in library prediction. Trypsin was set as the protease, with a maximum of 1 missed cleavage. Cysteine carbamidomethylation was enabled as a fixed modification. An FDR of 1% used for identification-level cut-offs. The DIA-NN output was loaded into Perseus version 1.6.10.50, and the matrix was filtered to remove all proteins identified as potential contaminants or decoy hits. LFQ intensities were log2(x)- transformed, and data were filtered to retain proteins with a minimum of two peptides and at least 3 valid LFQ intensities in one group. Data were normalised by subtracting column medians, and missing values were imputed from the normal distribution with a width of 0.3 and downshift of 1.8. To identify quantitatively enriched proteins between EV isolation methods, two-sided Student’s t-tests were performed with a permutation-based FDR of 0.05.

### Flow cytometry

Flow cytometry was performed to assess surface expression of the MISEV marker CD63. Samples were diluted in PBS and stained with CD63-PE (H5C6, Invitrogen; 12-0639-42) for 20 minutes prior to further dilution and analysis. A perfectly matched isotype control antibody (IgG1 κ -PE, Invitrogen; 12-4714-82) was used to determine background binding. For data collection a Becton Dickinson Accuri C6 flow cytometer was used with 2 lasers (488nm & 640nm), 4 detectors (533/30BP, 585/40BP, 670LP, and 675/25BP) and 5,000 events were collected. Analysis was carried on Floreada (https://floreada.io) and Flowjo (v10.10.0). Mean fluorescence intensities (MFI) and percentage positive data were generated by gating EVs on FSC-A/SSC-A.

### Cryogenic transmission electron microscopy

All samples were prepared for Cryo-TEM using an FEI Mark IV Vitrobot© held at 4°C and 100% humidity. For each of the three EV samples, 3.5 µl was loaded onto a plasma cleaned lacey carbon coated copper TEM grid (EM resolutions) before being blotted and then rapidly plunge frozen in liquid ethane. Transfer into the microscope was done using a Gatan-914 cryo-TEM holder, and the temperature was maintained below -165°C during analysis. All Cryo-TEM images were obtained using an FEI Titan3 Themis G2 equipped with a monochromator operating at 300 kV and fitted with a Gatan One-View CMOS camera.

### Enrichment analysis

Functional annotation and enrichment analysis of the proteins identified were performed using the ShinyGO v0.85 bioinformatics platform. Protein identifiers from each isolation method were mapped against the *Homo sapiens* database to retrieve Gene Ontology (GO) terms. Enrichment significance was determined using a hypergeometric distribution, with p-values adjusted for multiple testing using the False Discovery Rate (FDR). Terms were ranked primarily by Fold Enrichment (FE), defined as the ratio of the number of identified genes in a category to the number of genes. To facilitate a comparative analysis of the functional landscapes provided by each isolation platform, only the enriched proteins from each method were selected and cross-referenced to identify method-specific functional signatures.

## Results

### Workflow of the isolation methods

The experimental design for the comparative assessment of sEV isolation methods is illustrated in Figure 1. Conditioned media was subjected to the same pre-clearing steps (2,000 x*g* centrifugation followed by 0.22µm filtration) to remove cellular debris and large particles. The resulting filtered supernatant was then divided into three equal volumes for simultaneous processing by UC, SEC, and the EXODUS integrated separation system.

**Figure 1.**
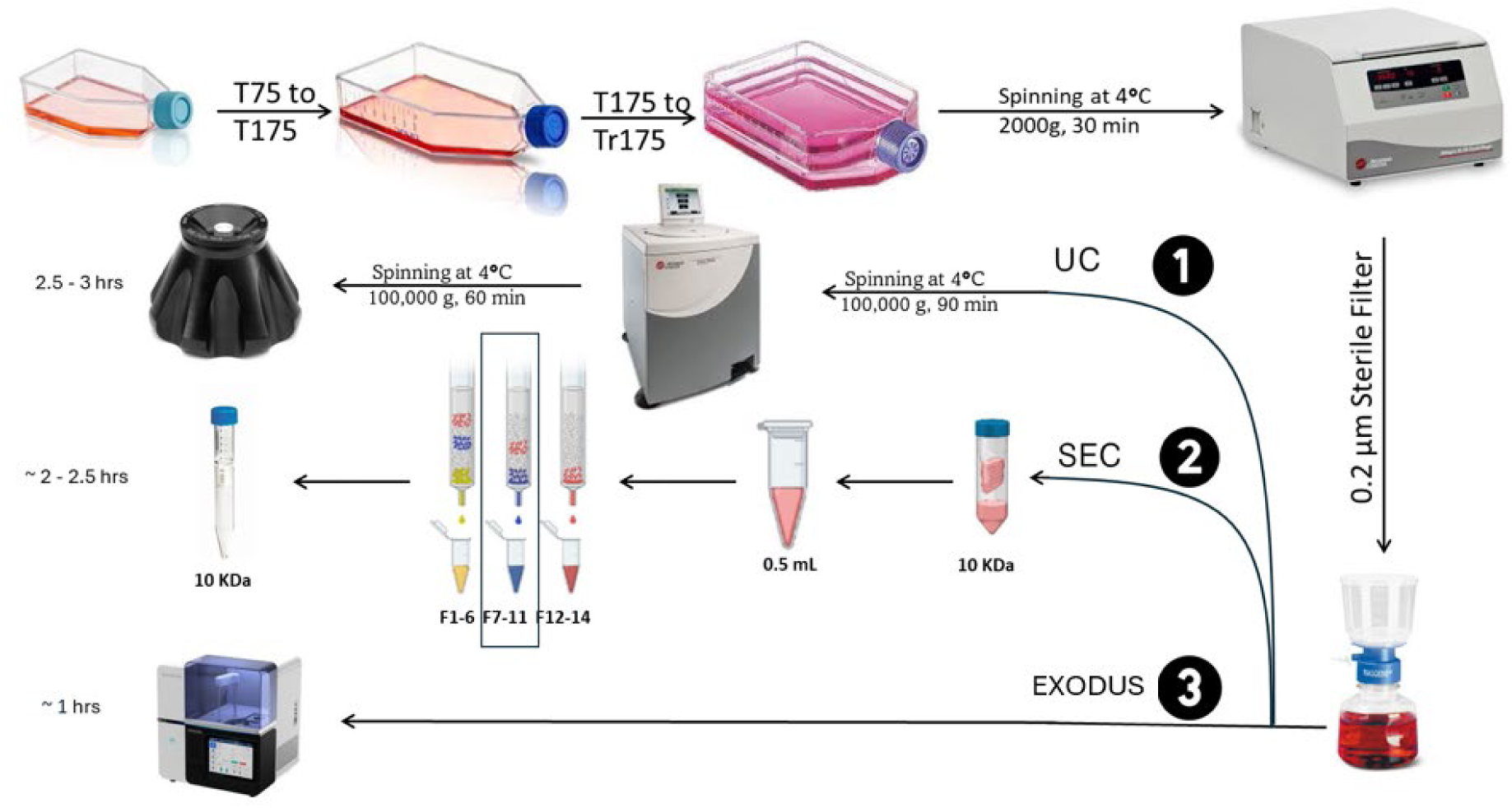
Schematic workflow for sEV Isolation from breast CAF conditioned medium: The comparative process involved sequential ultracentrifugation (UC), size-exclusion chromatography (SEC) to isolate a purified sEV fraction (F7-11), and EXODUS H600 system

A key differentiator among the methods was the time required for the final sEV isolation per sample. The automated EXODUS system required approximately 1 hour for completion. In contrast, the UC method, which requires two sequential 100,000 xg spins (90 minutes followed by a 60-minute wash), takes approximately 3 hours. The SEC workflow, which involved pre-concentration, column separation, collection of fractions 7–11 based on prior optimisation (Shelton *et al*., 2025), and subsequent re-concentration, required an estimated 2.5 hours.

### Independent NTA platforms confirm enhanced EV recovery with EXODUS and SEC relative to UC

To evaluate the efficiency of different sEV isolation strategies, samples were characterised using two independent NTA platforms: Nanosight and Zetaview. According to Nanosight analysis, EXODUS and SEC yielded comparable particle concentrations (1.43 ± 0.08) x 10^10^ particles/mL and (1.39 ± 0.2) × 10^10^ particles/mL, respectively, representing an approximate 6-fold increase in yield over UC (2.42 ± 0.4) × 10^9^ particles/mL) (Fig 2 A-C, Table 1). sEVs were also analysed using Zetaview. EVs isolated using EXODUS had a mean concentration of (5.57 ± 9.03) × 10^10^ particles/mL, representing a 25-fold increase over SEC (2.20 ± 0.62) × 10^9^ and a higher recovery improvement over UC, which yielded only (6.90 ± 3.92) × 10^7^ particles/mL (Fig 2 D-F, Table 1). Across both instruments, particle analysis of sEVs isolated using UC consistently yielded the lowest particle recovery, suggesting significant sample loss during the high-speed pelleting process. When combining data from both analytical platforms, overall particle recovery was significantly higher for EXODUS ((1.11 ± 0.48) × 10^10^ particles/mL) and SEC ((1.15 ± 0.47) × 10^10^ particles/mL) compared to UC ((0.151 ± 0.75) × 10^10^ particles/mL), representing an approximate 7-fold increase in yield (Figure 2G, Table 1). The physical dimensions of the isolated particles were analysed across both NTA machines. Both EXODUS and SEC methods produced populations with high similarity between the two NTA machines. On the Nanosight, EXODUS showed a mean diameter of 149 ± 5.7 (mode: 127 ± 3.95 nm), while SEC showed a mean of 143 ± 2.98 (mode: 129 ± 6.24 nm). Samples isolated using UC exhibited a shift toward larger particle sizes on the Nanosight platform with a mean diameter of 160 ± 2.66 nm (mode: 124 ± 0.38 nm). Interestingly, Zetaview analysis of UC samples yielded a smaller mean size of 118.4 nm ± 24 nm (mode: 103.6 ± 39), highlighting the influence of instrument-specific detection thresholds on the characterisation of EVs isolated using UC (Table 1) (Figure 2 H-I).

**Figure 2.**
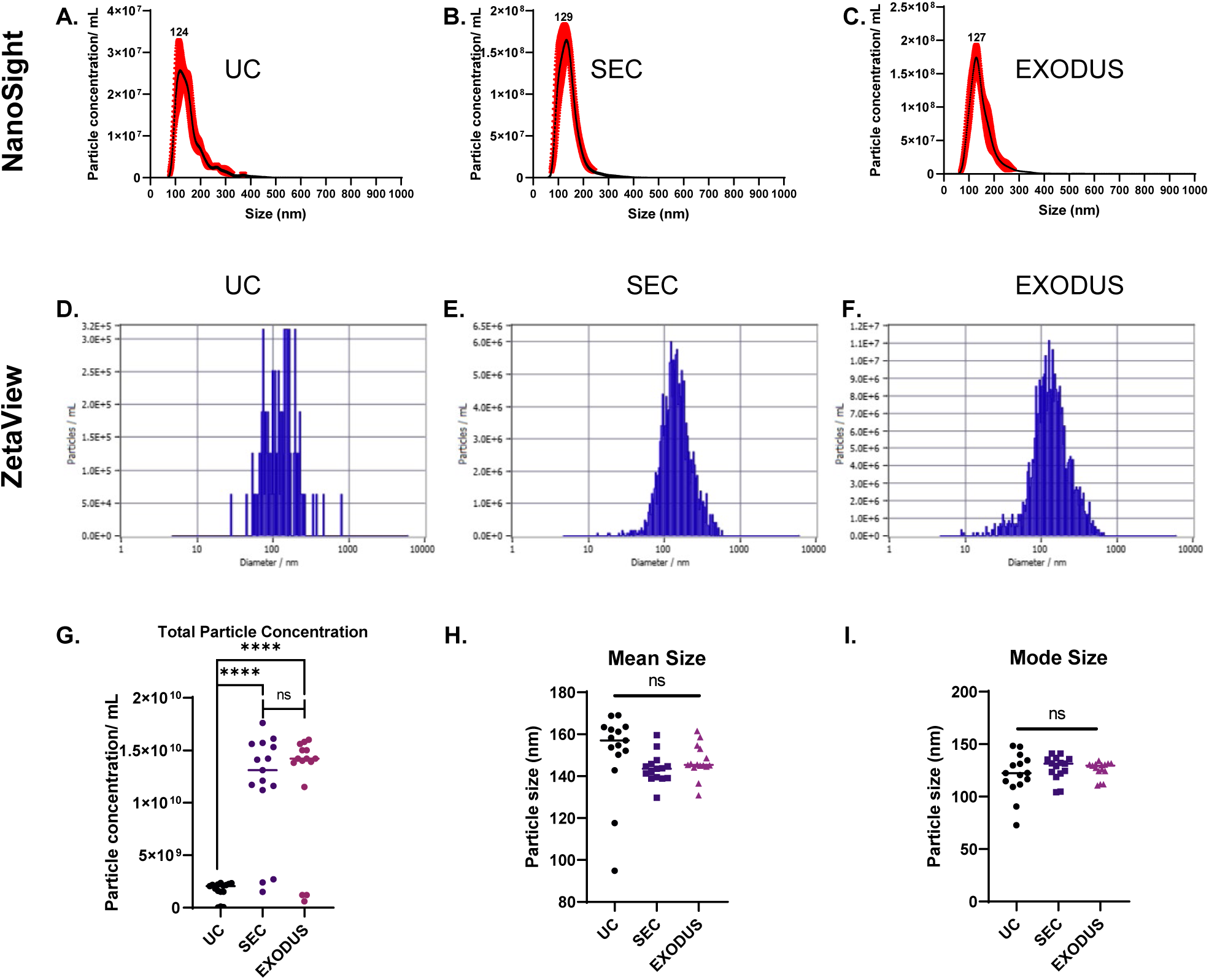
Particle characteristics of sEVs isolated by different methods: **(A-C)** Combined NTA (nanosight) profiles of each method showing the size distribution and enrichment of particles <200 nm (n=3). **(D-F) Representative images of the NTA profile using** ZetaView platform n= 3. **(G)** Total particle concentration identified by both NTA platforms **(H)** Mean particle size identified by both platforms and **(I)** mode particle size identified by both platforms. Statistical significance was assessed by one-way ANOVA followed by Tukey’s multiple comparisons test (****P<0.0001; ns, not significant). Data are mean ± SD.

**Table 1:**
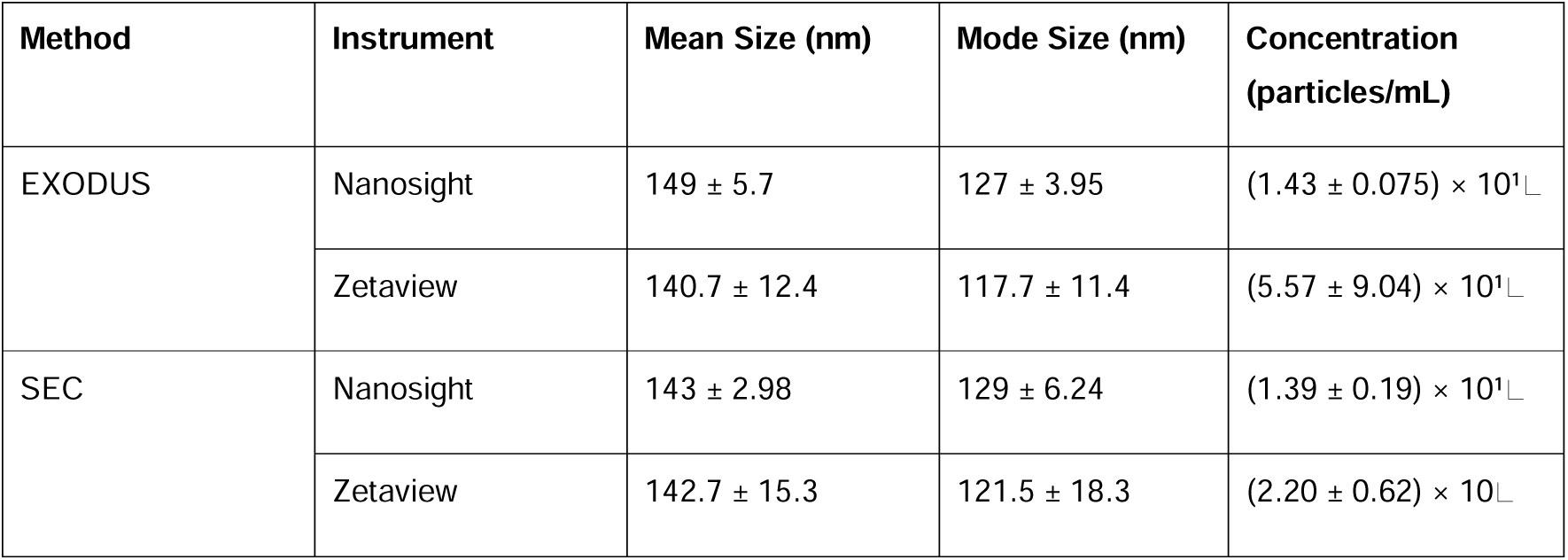
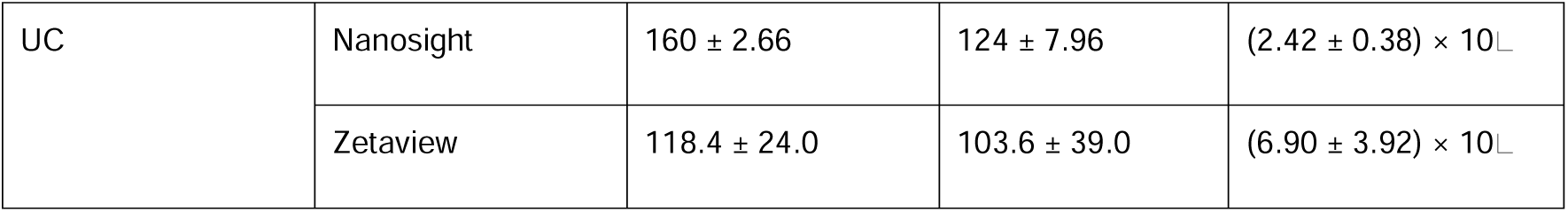
Physical characterisation of sEVs by Nanosight and Zetaview.

### Morphological analysis identifies distinct purity profiles and size distributions across EV isolation methods

To characterise the morphology, physical dimensions, and purity of the isolated EVs, Cryo-TEM was performed on all three preparations. All isolation methods yielded vesicles with characteristic intact lipid membranes and well-defined, rounded structures (Figure 3). Quantitative analysis of manually annotated particles (n = 150 per preparation) revealed distinct, method-dependent size profiles. EXODUS-isolated EVs exhibited the smallest and most uniform dimensions, with a median diameter of 66 nm, a mean of 75 nm, and a narrow, unimodal distribution peaking in the 40–80 nm range (Figure 3G, Table 2). In contrast, SEC-isolated EVs were slightly larger (median 74 nm, mean 93 nm); while their primary peak remained at 40–60 nm, the distribution was enriched for vesicles >80 nm, shifting the 75th percentile to 123 nm (Figure 3H, Table 2). UC-isolated EVs were the largest and most heterogeneous, with a median of 98 nm, a mean of 110 nm, and the widest size distribution (75th percentile = 139 nm) (Figure 3I, Table 2). Preparation background varied markedly across methods. EXODUS micrographs showed high clarity with no visible non-vesicular material (Figures 3A&D). SEC preparations contained an observable background of non-vesicular material (Figure 3B). UC preparations exhibited substantial amounts of co-isolated material (Figure 3C).

**Figure 3.**
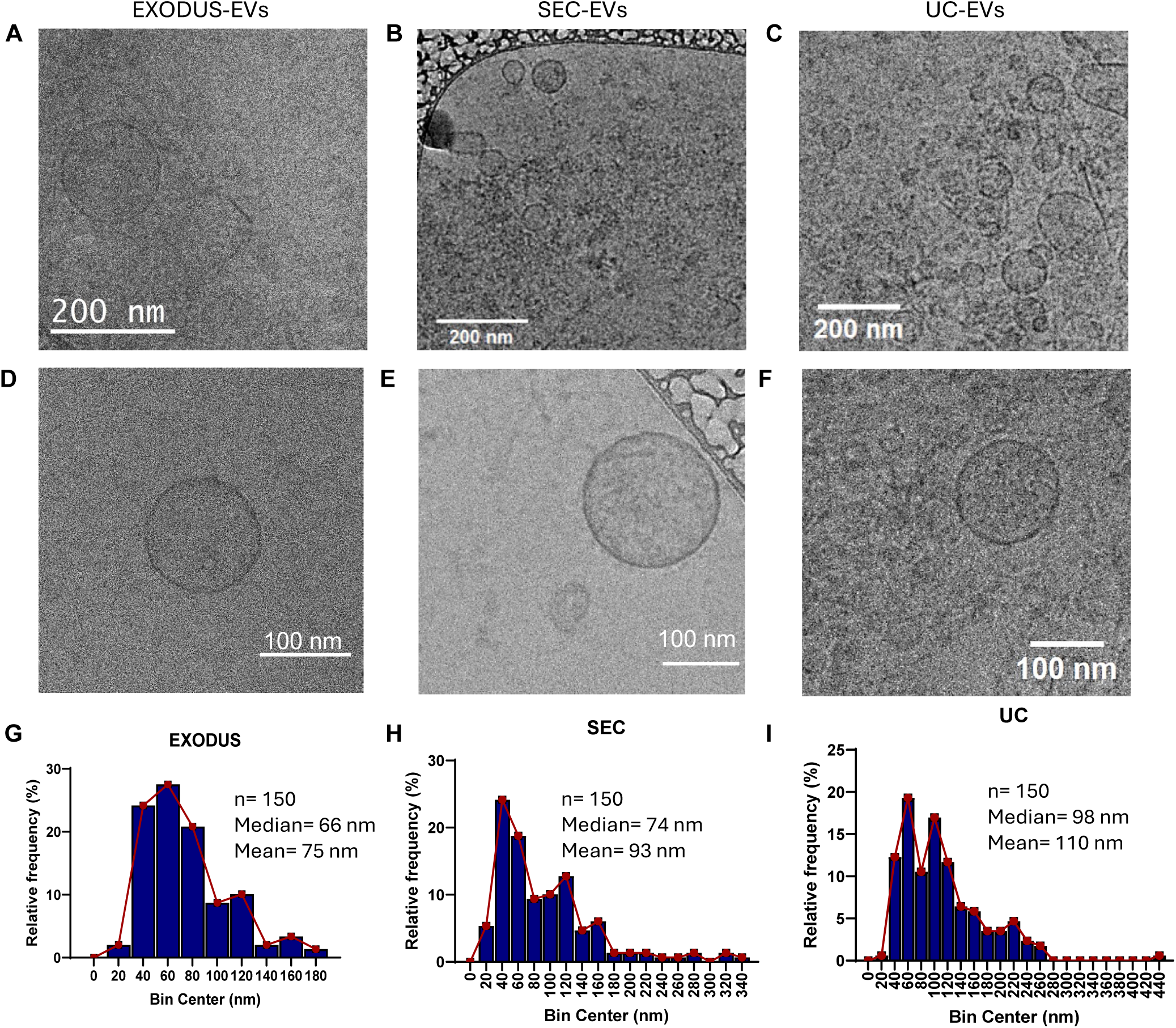
Morphological characterisation and size distribution of CAF-derived sEV using CryoEM: Representative cryo-electron microscopy (cryo-EM) images and corresponding size distribution profiles of sEVs isolated from B-CAF conditioned media. **(A–F)** Cryo-EM micrographs of sEVs isolated via **(A, D)** EXODUS nanofiltration, **(B, E)** size exclusion chromatography (SEC), and **(C, F)** ultracentrifugation (UC). **(A–C)** show wide-field views (scale bars = 200 nm), while **(D–F)** show high-magnification views of individual vesicles (scale bars = 100 nm). **(G–I)** Relative frequency histograms of vesicle diameters determined from cryo-EM images (n = 150 particles per method) for **(G)** EXODUS, **(H)** SEC, and **(I)** UC.

**Table 2:**
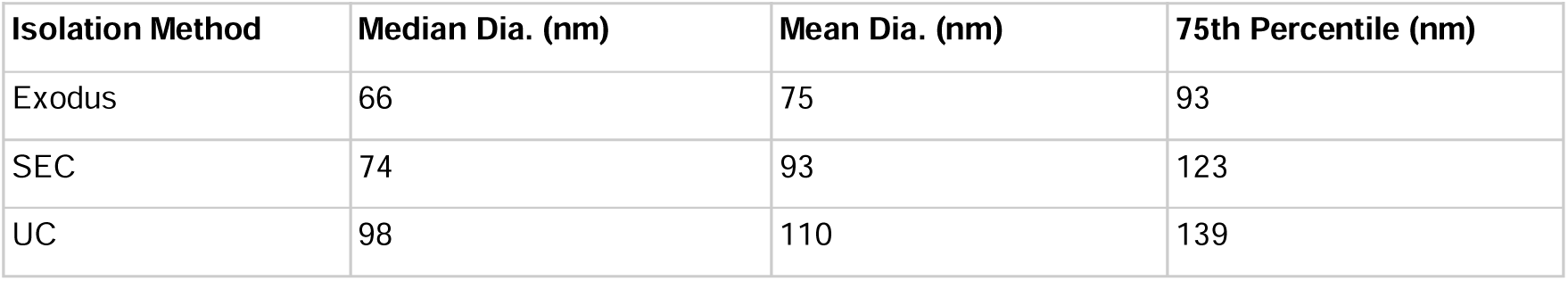
Summary of morphometric analysis across isolation methods.

### MISEV-compliant characterisation reveals method-dependent canonical EV protein signatures

EV preparations were characterised by Western blot for canonical transmembrane markers (CD63, CD9, CD81), cytosolic marker ALIX, and the ER-resident negative marker calnexin (CNX). All samples were normalised to a GFP spike-in added prior to microBCA quantification to ensure equivalent EV loading across preparations (Figure 4A). According to densitometric analysis, ALIX was most abundantly enriched in EXODUS preparations, with normalised band intensity approximately 1.25-fold greater than SEC (p<0.05) and substantially exceeding UC, where ALIX was detected at near-undetectable levels (p<0.001) (Figure 4B). CD81 followed a similar pattern, with EXODUS yielding significantly higher normalised intensity than both SEC (p<0.05) and UC (p<0.001), and SEC in turn significantly exceeding UC (p<0.01) (Figure 4C). CD9 was significantly enriched in EXODUS relative to UC (p<0.05), with SEC showing an intermediate level that did not reach statistical significance versus either method (Figure 4D). CD63 was detected at comparable levels across all three preparations with no statistically significant differences between methods (Figure 4E). Calnexin was not detected in all three EV preparations, confirming the absence of ER material consistent with MISEV2023 Category 4 negative marker requirements across all isolation strategies (Figure 4A). A notable difference was observed in the glycosylation status of CD63 and CD9. UC and SEC isolated sEVs exhibited both the lower molecular weight core protein form and the higher molecular weight glycosylated form of both tetraspanins (Figure 4A). In contrast, EXODUS-isolated EVs displayed the unglycosylated form exclusively, with the upper glycosylated bands completely absent (Figure 4A). To assess surface-expressed CD63 on intact vesicles, flow cytometric analysis was performed. Surface CD63 enrichment was significantly higher in EXODUS preparations compared to UC (approximately 19-fold vs 2-fold over IgG control; p<0.05), whilst SEC isolated-EVs showed an intermediate level of approximately 10-fold over IgG with no statistical significance relative to either method (Figure 4F).

**Figure 4.**
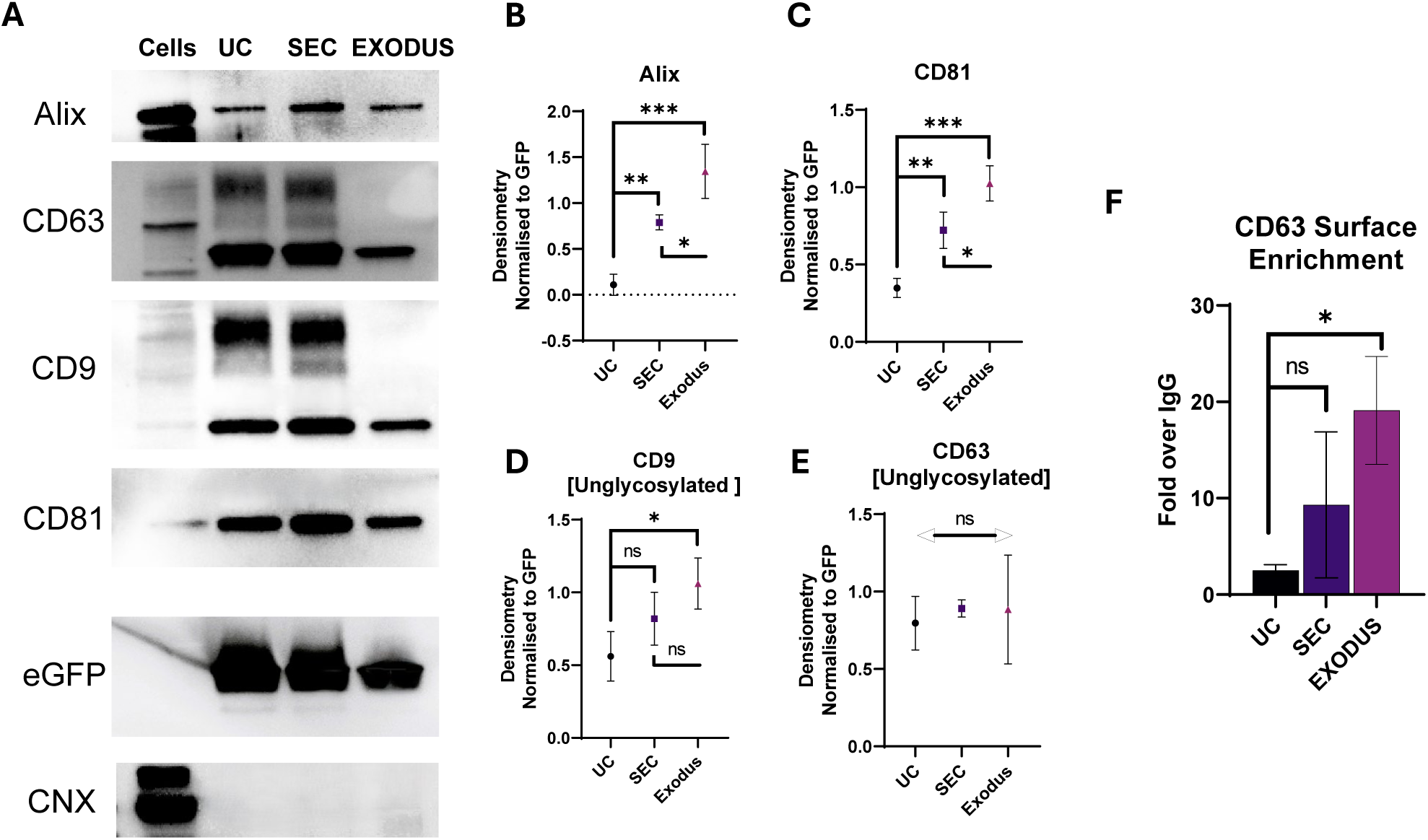
Comparative validation of EV marker enrichment and purity across isolation methods. **(A)** Western blot analysis of EV markers (Alix, CD81, CD9, CD63) and the negative control Calnexin (CNX) in parental cell lysates and EV fractions isolated by ultracentrifugation (UC), size-exclusion chromatography (SEC), and EXODUS. GFP serves as a loading control for EV lysates. **(B–E)** Densitometric quantification of **(B)** Alix, **(C)** CD81, **(D)** CD9, and **(E)** CD63. Western blot signals, normalised to GFP. Bars represent mean ± SD. Statistical significance was determined by one-way ANOVA with Tukey’s post-hoc test (*p < 0.05, **p < 0.01, ***p < 0.001), n=3 **(F)** Flow cytometric analysis of surface CD63 on sEVs. Data are expressed as MFI enrichment relative to the IgG isotype control (mean ± SD; *p < 0.05 vs. UC, n=3).

### Quantitative proteomic profiling of CAF-derived EVs across isolation methods reveals method-dependent subpopulation enrichment

DIA-based LC-MS/MS analysis identified 1289, 997, and 3172 proteins in EXODUS, SEC, and UC isolated B-CAF-derived sEV preparations, respectively (Supplementary file 1). Principal component analysis (PCA) of proteomic datasets showed 81.7% of total variance across PC1 (67.6%) and PC2 (14.1%), with all three methods forming tight, reproducible clusters that segregated clearly along both principal components, establishing isolation methodology as a primary determinant of sEV protein composition (Figure 5A). Statistical analysis of the quantitative data revealed substantial proteomic differences between all three pairwise comparisons at a 5% permutation-based FDR, with the greatest number of significantly different proteins observed between UC and both EXODUS (Figure 5B) and SEC (Figure 5C). The comparison between EXODUS and SEC revealed fewer differences, indicating greater proteomic similarity between the two methods (Figure 5D). A core proteome of 858 proteins was shared across all three methods, representing the core sEV population of B-CAF (Figure 5E). Notably, SEC-EVs demonstrated the highest degree of specificity, with 86.1% of detected proteins residing within the shared overlap. In contrast, UC yielded the largest unique proteome (1,829 proteins), of which only 27.1% were shared. This suggests that while UC provides a high-depth proteomic profile, it may be susceptible to significant co-isolation of non-vesicular proteins. EXODUS, on the other hand, occupied a middle ground, with 66.6% of its proteins shared with the other methods. The additional proteins identified by EXODUS compared to SEC may reflect a more comprehensive enrichment of membrane-associated cargo that is otherwise lost during passage through the SEC resin. However, further validation is required to distinguish them from potential co-isolates.

**Figure 5.**
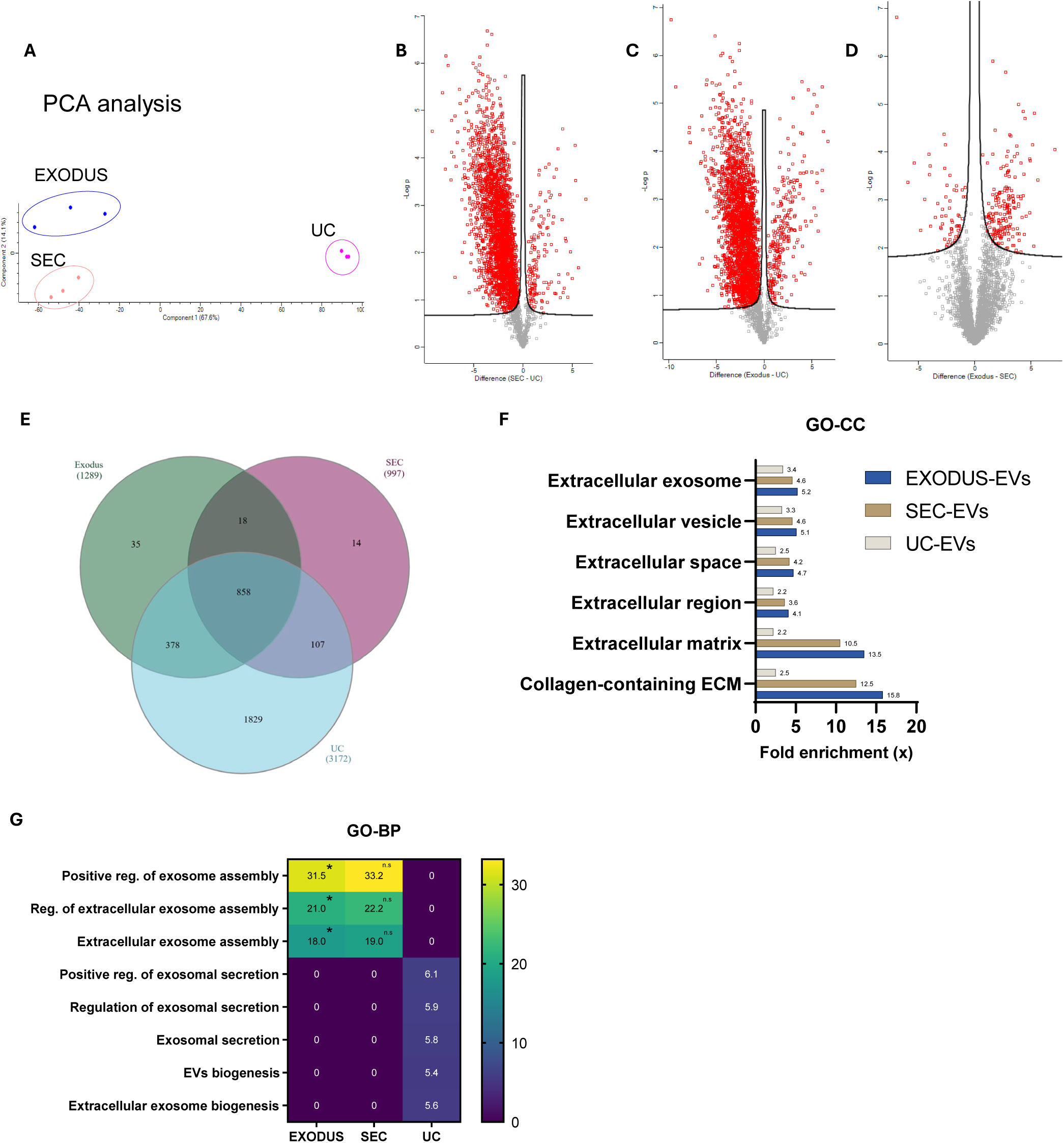
Proteomic characterisation and comparative analysis of EV isolation methods. **(A):** PCA of the proteomes obtained from EXODUS (blue), SEC (red), and UC (magenta) methods with distinct clustering by isolation method **(B-D)** Volcano plots highlighting proteins significantly enriched or depleted in **(B)** SEC vs. UC, **(C)** EXODUS vs. UC, and **(D)** EXODUS vs. SEC. Horizontal dashed lines indicate the threshold for significance (5% permutation-based false discovery rate, FDR). **(E)** Venn diagram of proteins identified across the three isolation methods. A core set of 858 proteins was common to all methods, with subsets unique to each. **(F)** Gene Ontology Cellular Component (GO-CC) enrichment analysis. Comparison of fold enrichment for EV-related terms EXODUS-EVs (blue), SEC-EVs (brown), and UC-EVs (grey). **(G)** Heatmap of Gene Ontology Biological Process (GO-BP) terms. The colour scale represents enrichment scores for BP. (*) indicated FDR score of <0.05. (n.s = non-significant).

### All methods satisfy MISEV2023 criteria but differ in preparation specificity and intracellular compartment profile

Assessment of the shared core proteome against the MISEV2023 protein reporting framework confirmed that all three isolation methods enriched for canonical transmembrane EV markers (CD9, CD81, CD63; MISEV Category 1) and cytosolic EV-associated proteins (TSG101, ALIX/PDCD6IP, Flotillin-1/2, Syntenin/SDCBP; MISEV Category 2), satisfying the minimum identity criteria for EV preparations. However, the methods diverged significantly in enrichment specificity and co-isolated material profiles beyond this shared core. Gene ontology cellular component (GO-CC) enrichment analysis of the proteins enriched in each method revealed that, despite its largest proteome, UC yielded the lowest quantitative enrichment for canonical EV terms, including extracellular exosome (FE = 3.4) and extracellular vesicle (FE = 3.3), reflecting dilution of true EV signal by co-sedimented non-vesicular material (Figure 5F, Supplementary file 2). SEC achieved intermediate enrichment for core EV terms (FE = 4.6 for both categories), while EXODUS demonstrated higher enrichment for extracellular exosome (FE = 5.2) and extracellular vesicle (FE = 5.1) (Figure 5F). EXODUS higher recovery for proteins associated with the extracellular matrix (FE = 13.5) and collagen-containing ECM (FE = 15.8), compared to SEC (FE = 10.5 and 12.5) and UC (FE = 2.2 and 2.5), respectively (Figure 5F). Furthermore, UC exhibited a high enrichment of non-vesicular background, uniquely showing significant enrichment for cytosolic ribosome (FE = 4.4), ER membrane (FE = 1.6), mitochondrion (FE = 1.2), all absent in SEC and EXODUS EVs (Supplementary file 2).

### Isolation method determines the functional maturity of the EV proteome landscape

To define the functional identity of the isolated particles, Gene Ontology Biological Process (GO-BP) analysis was performed (Supplementary file 3). EVs isolated via UC showed significant enrichment for terms associated with early-stage vesicle production and intracellular transport. Specifically, terms such as “extracellular vesicle biogenesis” and “extracellular exosome biogenesis” demonstrated fold enrichments of 5.5 and 5.6, respectively (Figure 5G). Notably, UC preparations lacked enrichment for terms related to the assembly of stable extracellular vesicles (Figure 5G). In contrast, EXODUS-derived preparations exhibited a more pronounced signature of assembled-EV-associated terms, including ’extracellular exosome assembly’ (FE = 18.0, FDR = 0.027), while notably lacking terms associated with the general, non-vesicular secretome (Figure 5G; Supplementary file 3). SEC preparations also identified the “extracellular exosome assembly” term with a high fold enrichment (FE = 19.0); however, this failed to reach statistical significance (FDR = 0.19) (Figure 5G). Furthermore, SEC identified only four secretion-related terms, none of which reached statistical significance (Supplementary file 2). Collectively, these data reveal a clear divergence in the molecular landscape captured by each method. The EXODUS platform selectively enriches for preparations with stronger assembled-EV-associated signatures while effectively excluding the soluble secretome and intracellular biogenesis machinery that characterise UC pellets. In this framework, SEC occupies an intermediate position, failing to reach statistical significance for either assembled vesicle signatures or general secretome terms (Supplementary file 3).

### Isolation method determines the matrisome landscape of CAF-derived EV preparations

To characterise the extracellular matrix (ECM) protein content of each preparation, enriched proteomic datasets from each isolation method were annotated against the NABA matrisome database and categorised into core matrisome and matrisome-associated divisions (Naba et al., 2012) (Figure 6A–D, Supplementary file 4). Matrisome proteins were detected across all three methods, though their total number and composition varied substantially. UC preparations had the highest number of matrisome proteins overall (n = 196), followed by EXODUS (n = 145) and SEC (n = 55) (Figure 6A). Within each preparation, core matrisome proteins, which encompass collagens, glycoproteins, and proteoglycans, accounted for 68 of 196 (34.7%) in UC, 84 of 145 (57.9%) in EXODUS, and 30 of 55 (54.5%) in SEC, with matrisome-associated proteins constituting the remaining fraction in each method (Figure 6A).

**Figure 6:**
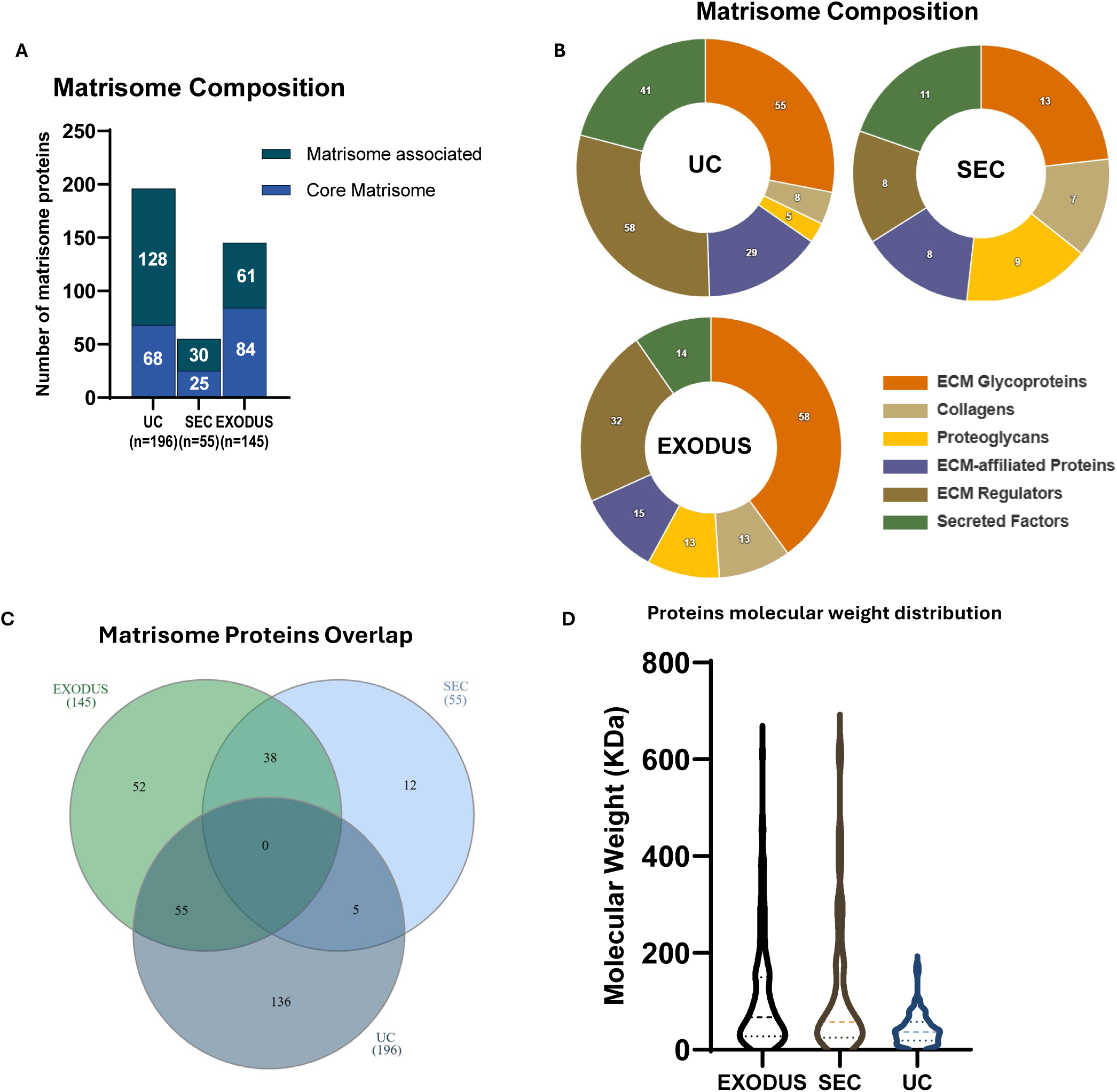
Matrisome landscape of CAF-derived EV preparations isolated using UC, SEC, and EXODUS. **(A)** Matrisome composition analysis of CAF-EVs categorised into core matrisome and matrisome-associated subtypes across the three isolation methods. Total number of identified proteins (n) is indicated for each method.(**B)** Donut plots showing the distribution of six distinct ECM-related protein classes identified within each isolation workflow. **(C)** venn diagram depicting the overlap of matrisome proteins across the three isolation methods. **(D)** Violin plots showing the molecular weight distribution (kDa) of the matrisome proteins identified in each preparation.

At the sub-category level, ECM glycoproteins represented the largest single category across all three methods, with 55 proteins in UC, 58 in EXODUS, and 13 in SEC (Figure 6B, Supplementary file 4). The EXODUS glycoprotein repertoire was broad, encompassing the full laminin heterotrimer complement (LAMA1–5, LAMB1–2, LAMC1), fibronectin (FN1), fibrillin-1 and 2 (FBN1, FBN2), all four thrombospondin family members (THBS1–4), fibulin1 and 2 (FBLN1, FBLN2), and tenascin-C (TNC). UC glycoproteins included fibronectin (FN1), thrombospondins 1–3 (THBS1–3), tenascin-C (TNC), periostin (POSTN), and a broad complement of IGFBP family members (IGFBP2–7), alongside multiple CCN family proteins (CCN1–3, CCN5). The SEC glycoprotein profile, while smaller in total number, included several large structural proteins not detected in UC, notably FRAS1, HMCN1, SVEP1, and three latent TGF-β binding proteins (LTBP1, LTBP2, LTBP4), which are involved in sequestration and extracellular presentation of TGF-β. Collagen-family proteins were higher in EXODUS (n = 13), followed by UC (n = 8) and SEC (n = 7). EXODUS enriched for additional collagen species absent from both other methods, including COL11A1, COL12A1, COL18A1, and COL5A2. UC collagen composition was characterised by the enrichment of COL6A1, COL6A2, and COL6A3, forming the type VI collagen heterotrimer, alongside COL4A1, COL8A1, COL13A1, COL14A1, and COL16A1. SEC and EXODUS shared a largely overlapping collagen repertoire (COL1A1, COL1A2, COL3A1, COL4A2, COL5A1, COL10A1, COL15A1), suggesting a common pool of fibrillar and network-forming collagen types. Proteoglycans followed a similar pattern, with EXODUS detecting 13 proteins, identifying large, aggregating proteoglycans and their stabilisers, such as aggrecan (ACAN), versican (VCAN), HAPLN1, and HAPLN3. SEC method-enriched proteins included seven proteoglycans with a profile containing small leucine-rich proteoglycans (SLRPs) such as biglycan (BGN), decorin (DCN), fibromodulin (FMOD), and lumican (LUM), none of which were detected in UC. The UC proteoglycan profile (n = 5) was restricted to basement-membrane associated factors like ESM1 and SPOCK1, suggesting that UC preferentially samples a soluble signalling niche rather than the structural matrisome.

UC preparations were distinguished by a substantially higher number of ECM regulators (n = 58), ECM-affiliated proteins (n = 29), and secreted factors (n = 41), compared to EXODUS (32, 15, and 14, respectively) and SEC (8, 8, and 11). Within the ECM regulators category, UC contained an expanded MMP repertoire (MMP1–3, MMP9, MMP11, MMP12, MMP14, MMP19), the full lysyl oxidase family (LOX, LOXL1–4), collagen prolyl hydroxylases (P4HA1, P4HA2, PLOD2), multiple cathepsins (CTSA, CTSB, CTSK, CTSL, CTSV, CTSZ), and the protease inhibitors TIMP1 and TIMP3. UC secreted factors were predominantly growth factors and cytokines, including VEGFA, VEGFC, FGF2, FGF5, HGF, TGFB1, PDGFC, PDGFD, and multiple CXC chemokines (CXCL1, CXCL3, CXCL5, CXCL6, CXCL8, CXCL10), as well as CCN family members CCN1, CCN2, CCN3, and CCN5. EXODUS ECM regulators included ADAMTS7 and ADAMTS12, MMP11, MMP12, BMP1, HTRA1, HTRA3, PLOD1, and PLOD3. SEC ECM regulators were characterised by serine protease inhibitors and inter-alpha-trypsin inhibitor heavy chains (ITIH1, ITIH2, ITIH3), alongside A2M, MASP1, and ADAMTS13, suggesting a distinct protease-regulatory environment in SEC-isolated EVs. The SEC secreted factor profile was notably dominated by S100 calcium-binding proteins (S100A6, S100A7, S100A8, S100A9, S100A14), which were exclusive to SEC and not enriched in either UC or EXODUS preparations (Supplementary file 4).

Overlap analysis of matrisome proteins across methods revealed limited shared detection between isolation methods (Figure 6C). No proteins were identified jointly across all three methods. A total of 55 proteins were shared exclusively between EXODUS and UC, 38 between EXODUS and SEC, and five between SEC and UC. Method-exclusive matrisome proteins were mostly present in UC (n = 136), followed by EXODUS (n = 52) and SEC (n = 12). Analysis of the molecular weight distribution of matrisome proteins identified in each preparation showed distinct interquartile range (IQR) profiles between methods (Figure 6D). UC-isolated matrisome proteins had a lower median molecular weight (36.2 kDa; IQR 19.6–55.8 kDa; range 3.0–172.1 kDa), with 69.5% of proteins falling below 50 kDa and only 3.2% exceeding 100 kDa. In contrast, EXODUS matrisome proteins exhibited a broader distribution (median 66.8 kDa; IQR 28.6–149.5 kDa; range 1.6–611.0 kDa), with 36.6% exceeding 100 kDa. SEC matrisome proteins showed a comparable range profile (median 56.9 kDa; IQR 26.6–131.0 kDa; range 6.6–611.0 kDa), with 30.8% above 100 kDa and 46.2% below 50 kDa. Together, these findings suggest that isolation methods act as filters for the CAF secretome. While UC enriches for small, matrisome-associated signalling factors, EXODUS and SEC preferentially capture the high-molecular-weight architectural components of the extracellular matrix (Supplementary file 4).

## DISCUSSION

The choice of sEV isolation strategy is widely recognised as a primary determinant of both vesicle yield and molecular composition, with direct consequences for downstream biological interpretation and translational relevance (Welsh et al., 2024). Many EV studies compare biological conditions after applying a single isolation method. In this study, we performed a controlled, side-by-side comparison of UC, SEC, and the EXODUS platform for the isolation of sEVs from B-CAF conditioned media. Our findings suggest that these three methodologies do not simply recover a single, homogenous sEV pool with varying degrees of efficiency. Instead, each technique enriches for a biologically distinct population of the CAF secretome, characterised by unique yield kinetics, purity profiles, and molecular cargo repertoires.

Looking at the small particle recovery efficiency, UC produced approximately 7-fold fewer particles per millilitre than either EXODUS or SEC. The low particle recovery of UC is consistent with the well-characterised limitation of high-speed pelleting approaches in other studies in which UC had the lowest particle yield among several methods, including SEC and microfluidic Exodisc and ultrafiltration (Dong *et al*., 2020; Grenhas *et al*., 2024). The size distributions measured by Nanosight and Zetaview for SEC and EXODUS samples were comparable; however, UC samples showed discordance, consistent with previous reports showing that Nanosight and Zetaview can yield significantly different concentration and size measurements for biological EV preparations (Bachurski *et al*., 2019). This analytical discordance likely reflects the inherent heterogeneity of UC samples, where the presence of protein aggregates and non-vesicular components, along with instrument-specific detection thresholds, makes UC measurements particularly susceptible to background noise. While the cross-platform consistency between SEC and EXODUS size distributions might suggest mutual validation, such a conclusion must be approached with caution. Regardless of the NTA platform, NTA calculates the hydrodynamic radius of particles based on Brownian motion; therefore, both instruments share a fundamental limitation in that they enumerate total particles rather than confirmed vesicles and cannot definitively distinguish between lipid-bound structures and dense proteinaceous aggregates (Comfort *et al*., 2021; Kowkabany and Bao, 2024).

To contextualise the NTA findings, Cryo-TEM was employed to interrogate particle identity and assess sample purity at the single-vesicle level. Across all three isolation methods, Cryo-TEM size distributions were systematically shifted toward smaller diameters compared to NTA measurements. This well-documented phenomenon is attributable to two primary factors: first, orders of magnitude more particles than the ∼150 manually annotated by Cryo-TEM; and second, Cryo-TEM resolves the physical membrane-to-membrane diameter, whereas NTA measures the larger hydrodynamic shell (Bachurski *et al*., 2019; Noble *et al*., 2020). Crucially, Cryo-TEM provided definitive evidence of non-vesicular contaminants in both SEC and UC preparations, with UC exhibiting the highest degree of background interference, as frequently reported in the literature (Van Deun *et al*., 2014). The near-total absence of non-vesicular background in EXODUS-isolated samples enables a higher degree of analytical clarity in downstream molecular profiling, ensuring that observed biological signatures are accurately and confidently attributed to the vesicular fraction.

Proteomic profiling reinforced this interpretation. Despite its lowest particle yield, UC produced the largest total number of proteins (3172 proteins), of which only 27.1% were shared with the other methods. This observation of high protein amount but low vesicle purity is explained by the pronounced enrichment of non-vesicular contaminants. GO-CC analysis identified significant enrichment of cytosolic ribosome, ER membrane, and mitochondrial proteins exclusively in UC preparations, with no such terms present or reaching significance in EXODUS or SEC samples. This pattern is consistent with previous proteomic studies demonstrating that UC co-sediments ribonucleoprotein complexes, protein aggregates, and organellar debris alongside genuine EVs (Zhou, McNamara and Dittmer, 2020; Soares *et al*., 2023). The detection of these compartment-specific proteins is therefore likely to reflect non-vesicular material rather than genuine EV cargo, and their inclusion risks misattributing non-vesicular biology to sEV-mediated processes. This interpretation is reinforced by the relatively low fold-enrichment for canonical EV GO-CC terms in UC preparations (extracellular exosome FE = 3.4; extracellular vesicle FE = 3.3). The low enrichment for mature vesicle assembly terms, contrasted with the high enrichment of biogenesis precursors, suggests that UC recovers a bulk secretome fraction. Rather than a purified population of fully formed EVs, UC captures a heterogeneous mixture of early-stage intermediates and non-assembled proteinaceous material. EVs isolated via EXODUS and SEC produced broadly similar particle concentrations, as assessed by NTA, and comparable size distributions across platforms for both methods. This comparability at the physical level may be due to both isolation methods being based on particle hydrodynamic radius rather than sedimentation velocity, thereby avoiding the high shear force inherent to UC (Chen *et al*., 2021; Visan *et al*., 2022). At the protein level, the two methods diverged substantially.

SEC was the most specific method, with 86.1% of its 997 detected proteins residing in the core shared proteome, reflecting a high-specificity preparation with minimal unique protein content. EXODUS, by contrast, identified 1289 proteins with 66.6% shared, accessing an additional 431 proteins not captured by SEC. These EXODUS-exclusive proteins were selectively enriched for ECM-associated GO terms, suggesting that the EXODUS platform captures a subpopulation of matrix-associated or membrane-tethered vesicular cargo that is excluded from the SEC resin matrix. This interpretation aligns with evidence that sEVs carry surface-bound ECM proteins that may be sterically or electrostatically retained on the resin used in SEC (Krušić Alić *et al*., 2022). GO-BP analysis showed that EXODUS was the only method to achieve statistical significance for the term “extracellular exosome assembly” and was the only method for which no secretome-related terms were returned. The exclusion of non-vesicular material is most likely driven by the architecture of the EXODUS platform, which employs a dual-membrane nanofiltration assembly integrated with periodic negative pressure and ultrasonic oscillations (Chen *et al*., 2021). The high-frequency oscillatory mechanism in EXODUS seems to facilitate the removal of soluble protein complexes and secretome-derived contaminants, which are typically co-precipitate or co-elute during passive filtration and density-based sedimentation methods (Brennan *et al*., 2020; Benayas *et al*., 2023). EVs isolated via SEC showed a numerically similar fold enrichment to EXODUS for GO ‘’exosome assembly’’ but failed to reach statistical significance. The lack of significance likely stems from reduced proteome depth in the SEC preparations, most likely due to co-isolated non-vesicular proteins that interfered with detection and ultimately reduced the statistical power of the enrichment analyses. Also, EVs isolated via SEC returned four secretion-related terms. However, none were statistically significant, placing it in an intermediate position between EXODUS and UC with respect to soluble secretome co-isolation and mature EVs retention. This hierarchy, EXODUS > SEC > UC in terms of mature vesicle specificity, is consistent with an emerging consensus that size-based methods generate cleaner vesicular preparations than pellet-based approaches (Monguió-Tortajada *et al*., 2019; Soares *et al*., 2023).

Another observation was the complete absence of glycosylated CD63 and CD9 in EXODUS preparations, in contrast to SEC- and UC-isolated EVs Both CD63 and CD9 are heavily N-linked glycosylated tetraspanins, and their glycosylation patterns are known to vary across intracellular compartments and EV subpopulations (Matsuda *et al*., 2020; Wang *et al*., 2023). This finding suggests that the three isolation methods recover vesicle populations with distinct tetraspanin molecular profiles. The exclusively unglycosylated profile observed in EXODUS preparations may indicate preferential enrichment of a sEV subpopulation containing less mature glycoforms before complete Golgi-mediated processing. Alternatively, the mechanical processing steps in the EXODUS workflow may remove labile N-linked glycans from the extracellular loops of tetraspanins during isolation, thereby generating an apparent hypoglycosylated profile that may not reflect a true biological difference. Although the precise mechanism remains unclear and requires further glycomic validation, these results provide qualitative evidence that the three isolation methods do not recover identical vesicle populations from the same conditioned media source. This observation was further supported by PCA analysis, which revealed three distinct and tightly clustered populations corresponding to each isolation method. Interestingly, while western blot analysis showed comparable levels of unglycosylated CD63 across methods, the flow cytometric analysis of surface CD63 revealed a significant 19-fold enrichment in EXODUS preparations compared to UC. This discrepancy suggests that although the amount of unglycosylated CD63 protein may be comparable across the three isolation methods, the EXODUS platform may enrich for EVs with a higher density of surface-exposed CD63. Alternatively, this observation may reflect the reduced presence of non-vesicular contaminants in EXODUS preparations. Cryo-TEM analysis showed that SEC and particularly UC samples contained higher levels of non-vesicular material, which may coat the EV surface and reduce antibody accessibility to intact, membrane-enclosed vesicles (Försönits *et al*., 2025).

Beyond general proteomic considerations, matrisome analysis further elucidated the functional role of these vesicles within the tumour stroma. CAFs are the primary architects and key modulators of the tumour stroma (Yang *et al*., 2023). Specifically, CAF-secreted EVs are increasingly recognised as key vectors of ECM remodelling in the tumour microenvironment, depositing structural glycoproteins, matrix metalloproteinases, and pro-fibrotic signals that collectively prime the stroma for tumour progression and metastasis (Dourado *et al*., 2019; Liu *et al*., 2023). In the EXODUS preparation, the detection of the full laminin heterotrimer complement, multiple fibrillin and thrombospondin paralogs, and a broad collagen repertoire was notable, as these are canonical components of the provisional and tumour-associated ECM and are not typically considered classical EV cargo (Halper, 2021; Welsh *et al*., 2024). Their enrichment in EXODUS may reflect genuine packaging of architectural ECM proteins within or on the surface of CAF-derived EVs, consistent with emerging evidence that mesenchymal cells can load EVs with pre-assembled or proteolytically processed ECM fragments during matrix remodelling (Peris-Torres and Rodríguez-Manzaneque, 2026). However, it cannot be excluded that some large ECM complexes co-isolate with EVs, particularly given the tendency of fibronectin, collagen fragments, and laminin to form high supramolecular assemblies (Mouw, Ou and Weaver, 2014). The absence of matrisome proteins shared across all three methods is particularly striking given that the starting material was identical and underscores the risk of treating any single method’s proteome as a complete or faithful representation of the CAF EV matrisome. A key distinction was that the UC method appeared less able to retain high molecular weight matrisome proteins, which were detected in EXODUS and SEC preparations. The UC proteome was instead characterised by a higher proportion of matrisome-associated proteins including growth factors, chemokines, IGFBP family members, and an expanded MMP repertoire relative to core matrisome proteins. The ratio of matrisome-associated to core matrisome proteins was highest in UC preparations, intermediate in SEC, and lowest in EXODUS, which was predominantly enriched for core matrisome components.

The SEC proteoglycan and glycoprotein profiles are particularly intriguing. The exclusive detection of SLRP family members (BGN, DCN, FMOD, LUM) in SEC may reflect their association with the EV surface glycocalyx, consistent with evidence that mesenchymal cells actively sort proteolytically processed ECM fragments and SLRPs into EVs during matrix remodelling (Li *et al*., 2024). In cancer and injury contexts, these EV-associated SLRPs, such as Decorin, can function as mobile pan-RTK inhibitors to modulate distant tissue environments (Salomäki *et al*., 2008). Equally, the SEC secreted factor repertoire was dominated by S100 calcium-binding proteins, which were not enriched in both UC and EXODUS, a finding of high relevance to CAF-mediated metastasis and therapy resistance (Friedman *et al*., 2020). Collectively, these findings argue against a purely hierarchical view of isolation method performance and instead support a model in which each approach samples a biochemically and physically distinct subspace of the CAF secretome. This has direct implications for studies aiming to attribute specific matrix-remodelling or pro-tumorigenic functions to CAF EVs, as the observed matrisome composition will differ substantially depending on the isolation workflow employed. Standardisation efforts in the EV field have thus far focused primarily on vesicle yield and purity benchmarks, but the present data suggest that matrisome composition may be an equally sensitive and biologically meaningful metric for method harmonisation, particularly in the context of mesenchymal and stromal cell biology, where ECM production is a defining functional output.

In summary, this study demonstrates that the selection of isolation method is not a neutral technical choice, but a fundamental determinant of the biological narrative derived from the breast CAF secretome. Notably, while all three isolation strategies satisfied the minimum MISEV2023 identity criteria, our results argue that compliance with these reporting standards should not be equated with equivalent preparation quality. The substantial disparities in protein enrichment, co-isolates profiles, and subpopulation representation documented here demonstrate that MISEV2023 compliance is a necessary baseline, but an insufficient criterion for establishing methodological equivalence. Our side-by-side comparison reveals that EXODUS, SEC, and UC enrich for biologically distinct fractions of the CAF secretome. UC enables broad proteome recovery but with a greater contribution from non-vesicular co-isolates, whereas SEC and EXODUS yield preparations with stronger EV-specific enrichment. Ultimately, these findings underscore the necessity for researchers to move beyond binary “pass/fail” frameworks and align their isolation strategy with their specific biological questions to ensure that observed effects are confidently attributed to the EV fraction.

## Author Contributions

**M.K.Eldahshoury**: Conceptualization, data curation, formal analysis, methodology, project administration, supervision, validation, visualization, writing – original draft, writing – review and editing. **E. Moss**: Formal analysis, investigation, methodology, validation, visualization, writing – review and editing. **J. Gillet-Woodley**: Formal analysis, investigation, methodology, validation, visualization, writing – review and editing. **M.Hindle**: formal analysis, investigation, methodology, validation, visualization, writing – review and editing. **M. Ilett**: formal analysis, investigation, methodology, validation, visualization, writing – review and editing. **M.Collins**: Conceptualization, formal analysis, data curation, investigation, methodology, validation, visualization, writing – review and editing. **J. R. Boyne**: Conceptualization, data curation, formal analysis, funding acquisition, methodology, project administration, resources, supervision, validation, writing – review and editing.

## Acknowledgements

This study is funded by Leeds Beckett University. E.Moss is funded by a doctoral fellowship from Leeds Beckett University. J. Gillet-Woodley is funded by British Skin Foundation studentship (009/S/23) awarded to JRB. We acknowledge funding from the Engineering and Physical Research Council (EPSRC), UK via project grant no. EP/X040992/1. We also thank the Leeds Electron microscopy and Spectroscopy (LEMAS) centre for use of their facilities for Cryo-TEM analysis. M Hindle is supported by the School of Health Small Grant Scheme at Leeds Beckett University (P34139). Instrumentation in the biOMICS Mass Spectrometry at The University of Sheffield was funded by a grant from the BBSRC (BB/Z515796/1).

## Supplementary Files

**Supplementary File 1. Quantitative proteomic dataset of CAF-derived EV preparations isolated using UC, SEC, and EXODUS.** Comprehensive DIA-based LC–MS/MS proteomic dataset of extracellular vesicle (EV) preparations isolated from breast cancer-associated fibroblast (B-CAF) conditioned media using ultracentrifugation (UC), size exclusion chromatography (SEC), and EXODUS nanofiltration. Proteins were filtered and analysed using DIA-NN and Perseus with a permutation-based FDR threshold of 5%.

**Supplementary File 2. Gene Ontology cellular component enrichment analysis of proteins identified in EV preparations isolated using UC, SEC, and EXODUS.** Gene Ontology Cellular Component (GO-CC) enrichment analysis of proteins enriched in extracellular vesicle preparations isolated by ultracentrifugation (UC), size exclusion chromatography (SEC), and EXODUS. The dataset includes enriched GO terms, fold enrichment values, false discovery rate (FDR)-adjusted p-values, gene counts, and associated proteins identified using ShinyGO analysis against the Homo sapiens database.

**Supplementary File 3. Gene Ontology biological process enrichment analysis of proteins identified in EV preparations isolated using UC, SEC, and EXODUS.** Gene Ontology Biological Process (GO-BP) enrichment analysis of extracellular vesicle-associated proteins isolated from B-CAF conditioned media using ultracentrifugation (UC), size exclusion chromatography (SEC), and EXODUS. The file contains significantly enriched biological processes, fold enrichment scores, adjusted false discovery rate (FDR) values, gene counts, and associated protein lists.

**Supplementary File 4. Matrisome annotation and comparative extracellular matrix profiling of CAF-derived EV preparations.** Comprehensive matrisome annotation of proteins identified in extracellular vesicle preparations isolated using ultracentrifugation (UC), size exclusion chromatography (SEC), and EXODUS. Proteins were classified according to the NABA matrisome framework into core matrisome categories (collagens, ECM glycoproteins, proteoglycans) and matrisome-associated divisions (ECM regulators, ECM-affiliated proteins, and secreted factors).

